# Deep learning approach for automated cancer detection and tumor proportion score estimation of PD-L1 expression in lung adenocarcinoma

**DOI:** 10.1101/2020.05.31.126797

**Authors:** J. Wu, C. Liu, X. Liu, W. Sun, L. Li, Y. Zhang, J. Zhang, H. Wang, X. Liu, X. Yang, X. Huang, D. Lin, S. Ling

## Abstract

**Background:** This study proposed a computational method to detect the cancer areas and calculate the tumor proportion score (TPS) of PD-L1 immunohistochemistry (IHC) expression for lung adenocarcinoma based on deep learning and transfer learning.

**Patients and methods:** PD-L1 22C3 and SP142 IHC slides of lung adenocarcinoma samples on digitized whole-slide images (WSI) database were employed. We build a deep convolutional neural network (DCNN) to automatically segment cancer regions. TPS was calculated based on segmented areas and then compared with the interpretations of pathologists.

**Results:** We trained a DCNN model based on 22C3 dataset and fine-tuned it with SP142 dataset. We obtain a robust performance on cancer region detection on both datasets, with a sensitivity of 93.36% (22C3) and 92.80% (SP142) and a specificity of 93.97% (22C3) and 89.25% (SP142). With all the coefficient of determinations larger than 0.9 and Fleiss’ and Cohen’s Kappa larger than 0.8 (between mean or median of pathologists and TPS calculated by our method), we also found out the strong correlation between the TPS estimated by our computational method and estimation from multiple pathologists’ interpretations of 22C3 and SP142 respectively.

**Conclusion:** We provide an AI method to efficiently predict cancer region and calculate TPS in PD-L1 IHC slide of lung adenocarcinoma on two different antibodies. It demonstrates the potential of using deep learning methods to conveniently access PD-L1 IHC status. In the future, we will further validate the AI tool for automated scoring PD-L1 in large volume samples.

## 1. Introduction

Programmed cell death-1 (PD-1)/programmed cell death-ligand 1 (PD-L1) pathway-targeted immunotherapy has been proved as an important management of non-small cell lung cancer (NSCLC)^1–3^. This development prompted companion diagnostic immunohistochemistry (IHC) tests assessing the PD-L1 protein expression as a biomarker that predicts response to immunotherapy^2–5^. Different antibodies’ properties, expression quantitation, intra-tumoral heterogeneity and diverse spatial distribution patterns of different cell types^6–9^ have already brought challenges for PD-L1 IHC staining’s applications. Pathologists have difficulties in interpreting an accurate PD-L1 status on cancer sections which is a time-consuming clinical practice procedure. Additionally, inter-observer variation among pathologists may contribute to inaccurate patient stratification and limit the applications of PD-L1 IHC staining as an effective biomarker in clinical procedure^10–13^.

Advances in digital pathology and widely available scanner have set the stage for the clinical application of machine learning (ML), and make it possible to develop an assistive tools of Artificial Intelligences (AI) to improve the pathologic practice^14–16^. Deep convolutional neural network (DCNN) is specialized in digital image analysis tasks, and has been successfully applied to diagnostic pathology, including histopathological diagnosis^17^, cancer detection^18^, cell classification and enumeration^19–20^, mutation and microsatellite instability prediction^21–23^, tumor grading^21^, and cancer prognostication^22^. Utility of computer-assisted diagnostics showed the improvement of quantitative and qualitative pathologic interpretation of IHC staining, such as algorithms for IHC scoring of HER2, indicating the enormous potential of AI in assisting the pathologist with objective IHC scoring for stratified medicine^23^. Digital pathology technique may show advantages at the quantification analysis of PD-L1 IHC status and reproducibility and accuracy for interpretations of PD-L1 IHC scoring in a combination with AI and deep learning algorithms, as demonstrated in recent studies^24–26^.

In this study, we proposed a DCNN-based computational method to predict cancer region and calculate tumor proportion score (TPS) of PD-L1 to address those challenges focusing on lung adenocarcinoma, which has accounted for the largest proportion of patients with lung cancers in China and shows much more difficulties in evaluating PD-L1 expression due to its complex morphological patterns^28^. The aim of our study was to set up a generalized model for two different antibodies and explore the possibility to build a uniform strategy for antibodies with distinct characteristics.

## 2. Material and Methods

### 2.1 Study design

Two different antibodies were involved in this study. Firstly, we chose DAKO PD-L1 22C3 PharmDx assay to build up the standard model for cancer area detection and calculate tumor proportion score. SP142 was chosen to tune and generalize a new model to expand its application. Finally, we achieved a high performance on both 22C3 and SP142 dataset for cancer area prediction and TPS calculation. The workflow was shown in Fig.1.

**Fig.1.**
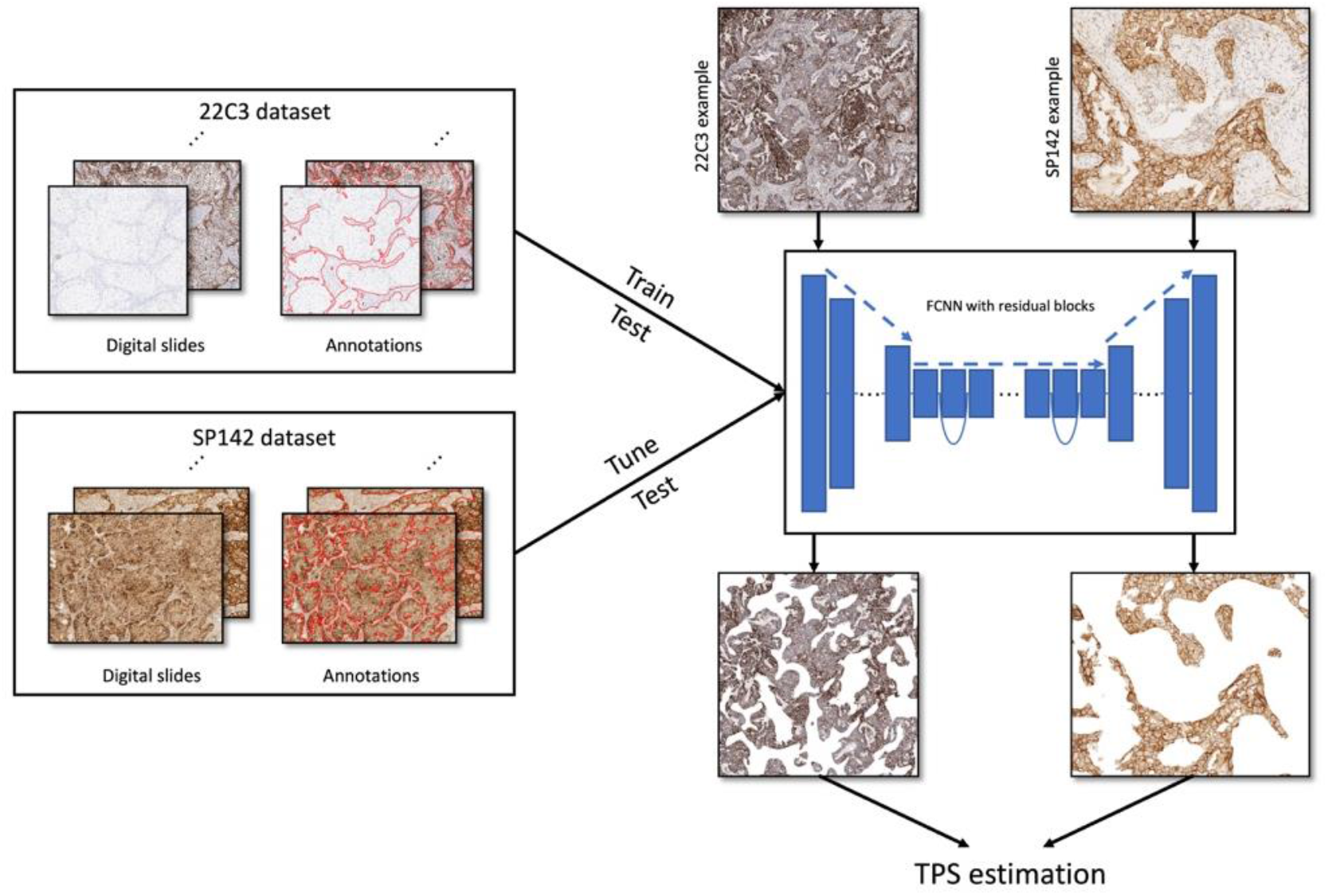
A general workflow of automated lung adenocarcinoma detection and automated TPS estimated in this study with 22C3 dataset to train and test the model, and SP142 dataset to tune the model.

### 2.2 Dataset

In this study, 115 slides from 115 patients were used. The information of patients is listed in Table.S1. All samples analyzed are unselected series from a collection of surgically resected lung adenocarcinomas specimens at Peking University Cancer Hospital & Institute from January 2017 to July 2018. This retrospective archival material and results did not have any impact on the management of patients. The Institute Review Board of Peking University Cancer Hospital & Institute, Key laboratory of Carcinogenesis and Translational Research (Ministry of Education) approved the study protocol.

All the slides were obtained by cutting formalin-fixed, paraffin embedded lung adenocarcinoma samples into 4 μm-thick sections. 50 slides were stained with PD-L1 IHC 22C3 pharmDx assay according to a standard staining protocol using appropriate automated staining devices (Dako Autostainer Link 48 platform). 65 slides were stained with anti-human PD-L1 rabbit monoclonal antibody (clone SP142, ZSGB-BIO, Beijing, China) at a working solution and incubated for 15 min at 37 °C on an automated staining platform (BOND-MAX, LEICA, Leica Biosystems Newcastle Ltd., Newcastle, UK).

All the slides were scanned by Leica Aperio CS2. We selected 3-5 cancer or stroma regions (>1mm^2^) on each slide (171 regions of 22C3 and 178 regions of SP142). In total 224 cancer areas and 123 stroma areas were selected for model building.

### 2.3 Manual annotations and tumor proportion scoring estimation

All the cancer and stroma area in selected regions were annotated by five pathologists for 22C3 samples and two pathologists for SP142.

Pathologists also needed to estimate the TPS for both datasets. TPS was calculated as the percentage of survival cancer cells exhibiting partial or complete density of membranous staining in NSCLC according to DAKO PD-L1 IHC 22C3 pharmDx Interpretation Manual, as the equation below.

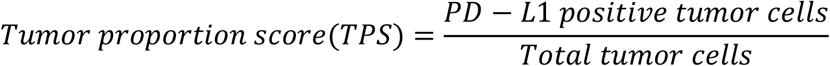

Pathologists’ scores were recorded on a 14-point scale representing PD-L1 expression as follows: negative or <1%, 1-4%, 5-9%, 10-19%, 20-29%,… 90-94%, 95-98%, 99-100%, with a range of 10% between 10% and 90% but narrower for both sides. Median and average TPS of 22C3 assay by five pathologists were taken as a final ground truth (GT). TPS of SP142 assay was calculated by two pathologists. The inconsistent cases were reviewed by a third consultant pathologist and a consensus of GT was taken subsequently. Time consumed for the whole process was recorded. Before this study, all pathologists involved were trained in scoring 22C3 assays by protocol.

### 2.4 Automated cancer area detection and tumor proportion scoring calculation

We first built a model based on 22C3 dataset for cancer area segmentation and test its performance including precision, sensitivity, specificity, F1-score, IoU and accuracy. 22C3 dataset was separated into a training set of 40 slides (130 regions, 8467 patches) and a test set of 9 slides, 41 regions (one slide was excluded due to poor quality of staining). We also test this model on SP142 dataset which contains 178 regions from 65 slides. After that SP142 dataset, containing a training set of 103 regions, 7008 patches and a test set of 75 regions, 6655 patches, was used to fine-tune the model and evaluate the performance. There are 23 regions without any cancer cells in the training set, which helped us get a conservative model. The workflow with details of dataset was shown in Fig.S1.

After cancer areas were detected by our model, we then segmented the cancer cells with watershed algorithm using QuPath^29^ (Version 0.1.2). We used a negative control group of 5 slides to determine the 3,3’-Diaminobenzidine (DAB) staining threshold. We defined pTPS for TPS estimated by pathologists and mTPS calculated by proposed method and will be used and compared below.

### 2.5 Statistical analysis

Data from the experts were compared with calculated scores by our method and the Kappa (Cohen’s and Fleiss’) parameters were calculated to validate the agreement using Python (Version 3.6). For 22C3, Fleiss’ Kappa was calculated for interpathologist’s agreement, but Cohen’s Kappa was calculated between the median and average data of pathologists and the score by proposed method. For SP142 Cohen’s Kappa was calculated for both inter-pathologist’s agreement and AI-pathologist’s agreement. We chose 1% and 50% as different cutoffs to assess the consistency between pathologists and AI method.

## 3. Results

### 3.1 Model training and performance testing

A pilot study was firstly performed to find the concordance of pathologists’ labeling. We calculated the intersection over union (IoU) and a high concordance between 3 random pathologists on 3 random samples and showed the results in Table.S2.

The performance of models trained with individual 22C3 dataset and fine-tuned with SP142 dataset was shown in Table.1. Evaluation on test set of 22C3 dataset showed a moderate sensitivity (72.86%) with a high specificity (93.51%), which is the performance of High-Specificity Model in Table.1. We trained the model on the same training set with different loss function and sample weights and get a balanced model with a sensitivity of 93.36% and specificity of 93.97% as shown in Balanced Model of Table.1. Based on the performance, we could conclude that our model for cancer area segmentation is robust enough (most parameters could reach a number over 90%) and it indicates the possibility to be applied on other antibodies.

**Table.1.** Performance of cancer area detection model. We calculated several parameters (precision, recall, specificity, F1-score, IoU and accuracy) to determine the performance of first model (trained with training set of 22C3 dataset) on test set of 22C3 dataset and SP142 dataset, and second model (tuned with training set of SP142 dataset) on test set of SP142 dataset.

|  |  | **Precision** | **Sensitivity** | **Specificity** | **F1-score** | **IoU** | **Accuracy** |
| --- | --- | --- | --- | --- | --- | --- | --- |
| **High-specificity model** | 22C3 | 92.84% | 74.79% | 80.15% | 70.36% | 91.08% | 96.19% |
|  | SP142 | 81.19% | 39.13% | 96.84% | 43.12% | 33.15% | 88.26% |
|  | SP142 tuned | 91.69% | 61.82% | 70.64% | 60.54% | 88.26% | 99.17% |
| **Balanced model** | 22C3 | 90.09% | 93.36% | 93.97% | 91.15% | 84.49% | 94.00% |
|  | SP142 | 83.54% | 86.26% | 86.24% | 82.63% | 74.15% | 88.52% |
|  | SP142 tuned | 90.63% | 92.80% | 89.25% | 91.22% | 84.34% | 92.80% |

### 3.2 Tumor proportion score of PD-L1 staining

Comparison of pTPS is shown in Fig.2(a) and a high consistency with each other was found in most of the cases for 22C3 dataset.

**Fig.2.**
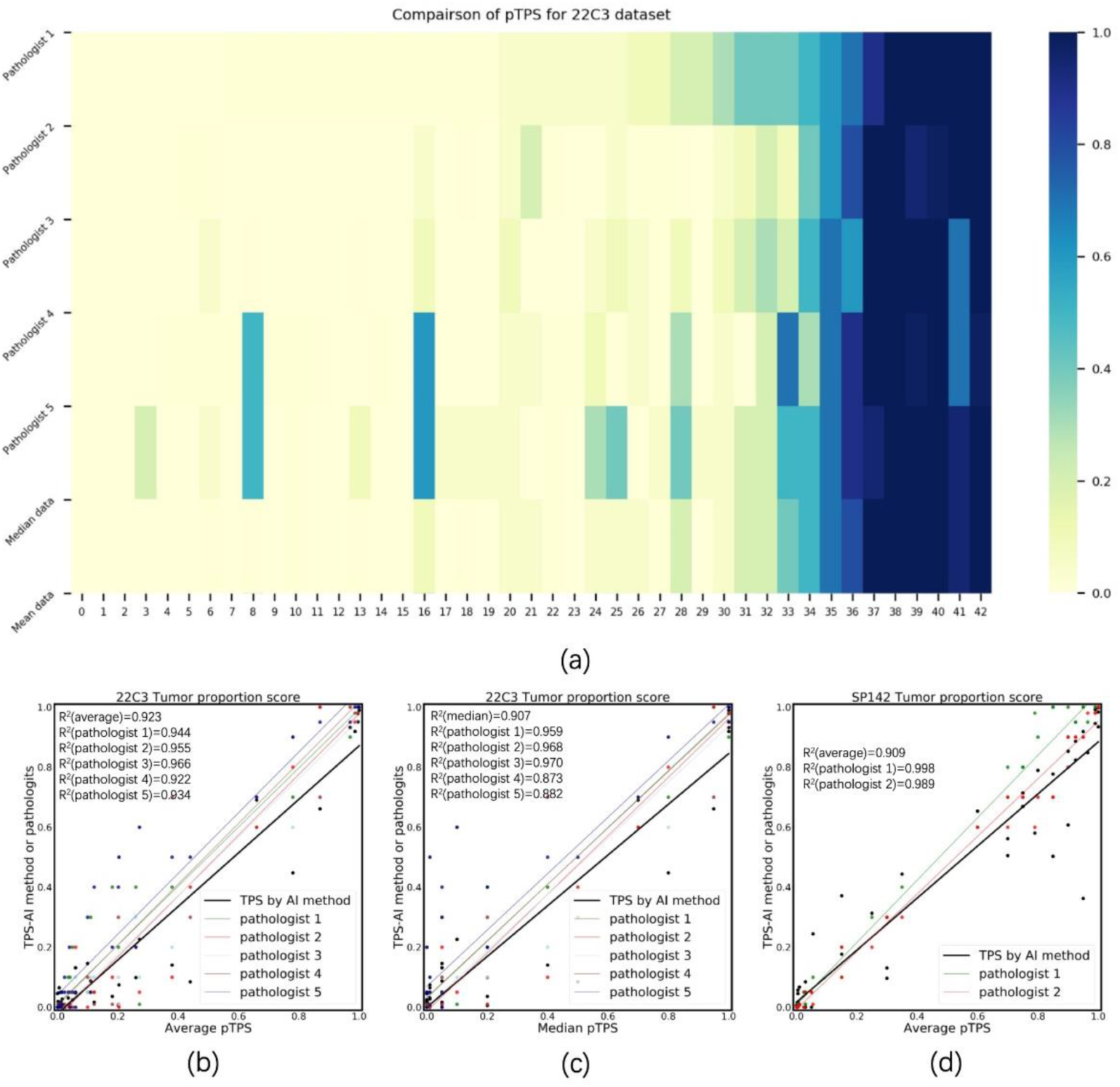
(a) Comparison of interpreted TPS from five pathologists for 22C3 dataset including the median and mean data. (b) Agreement of mean data of five pathologists and AI model for 22C3 dataset; (c) Agreement of median data of five pathologists and AI model for 22C3 dataset; (d) Agreement of mean data of two pathologists and AI model for SP142 dataset;

The model was applied on the 22C3 test set and SP142 test set to segment the cancer areas and calculate the mTPS based on them. We then compared the pTPS and mTPS and tried to find out the correlations between them. A highly significant correlation was found between average value of pathologists’ interpretations and automated PD-L1 analysis for both 22C3 dataset (R^2^=0.9234, p<0.0001) and SP142 dataset (R^2^=0.9098, p<0.0001), and also the median data from pathologists and AI model for 22C3 dataset (R^2^=0.9068, p<0.0001). Results were shown in Fig.2(b-d).

When TPS was categorized using cutoff values of 1% and 50%, the Kappa parameter was calculated to test the consistency between multiple results in Table.S3. Results showed a high consistency with each other when the cutoff was 50% with a Kappa value from 0.8 to even close to 1 (Kappa range from 0.841-1.000). However, the results from AI model and pathologists don’t show high consistency when the cutoff was set at 1% (Kappa range from 0.302-0.557)

### 3.3 Comparison of processing time for pathologists and AI model

Average time of review per region was obviously shorter with AI tool than manual performance which was shown in Table.S4. The average time reduced to 1.5 min/region if we run the proposed method on NVIDIA GeForce GTX 1080Ti to interpret the mTPS score, comparing with over 2 min/region for a pathologist (range 2.08-3.12 min/region for 22C3 and 2.98-.3.32 min/region for SP142) which varies based on their experience and experts.

## 4. Discussion

In this study, we endeavored to develop a deep learning method that enables cancer region detection and effectively scoring TPS of PD-L1 expression with a concrete value on PD-L1 IHC slides of DAKO 22C3 and SP142.

In recent years, it has been demonstrated that AI and deep learning-based methods could solve complex tasks in digital pathology image analysis of IHC profile to improve the reproducibility and accuracy^15^. However, previous studies didn’t make their effort to set up a generalized model or strategy to cover multiple antibodies, which could be accomplished with transfer learning. Transfer learning is a new frequently-used method to cover characteristics from different objects in one single model^27^. The model was trained based on the pre-trained model and could combine the properties of different dataset together. It provides us the access to build up a robust model for antibodies with distinct characteristics.

In our study, we first built a model for cancer area segmentation on 22C3 dataset and tested its performance. According to the Blueprint PD-L1 IHC Assay Comparison Project, SP142 showed largest inconsistency with other tested antibodies due to its poor tumor-specificity^7^. The clone SP142 in our study was even poorer in tumor specificity compared with Ventana, which was used to explore the generalization ability of our model trained by 22C3 dataset. What’s more, we applied the transfer learning^27^ to combine characteristics from different antibodies and increase the robustness further. A similar performance was obtained for both SP142 and 22C3 dataset by transfer learning model, confirming that the model could be interchangeable among antibodies by a small addition on training set.

There are still some limitations in our study. First of all, the pTPS results for WSI with very low percentage of positive cancer cells didn’t reach a high consistency among pathologists, and the current AI system seems to be less capable for this type of images, which led to a relatively low concordance between pathologists and AI method. The scattered immune cells which may show PD-L1 positive and intermixed in tumor nests cannot be easily recognized both by AI and pathologists. Last but not the least, although an excellent agreement between the values of TPS by experts and proposed method was achieved, it doesn’t mean that our current AI method can absolutely replace manual diagnosis. Due to limitation of training dataset, the model could detect all the cancer types in complex clinical practice, especially for rare lung cancer types. In the future, a multi-institutional project might be built up to co-generate a more robust model for estimation.

In conclusion, we proposed a deep learning model for automated cancer cell detection and assessment of PD-L1 expression across 22C3 and SP142 antibody in lung adenocarcinoma. It proves that advances in AI tools for immune biomarker researches are useful for cancer immunotherapy and might be an effective way to address the challenges associated with PD-L1 assessment in future clinical routines.

## 5. Acknowledgments

This work was supported by the National Natural Science Foundation of China (Grant No. 81871860).

## Reference

1. Garon EB, Rizvi NA, Hui R, et al. Pembrolizumab for the treatment of non-small-cell lung cancer. N Engl J Med 2015;372:2018–2028.

2. Borghaei H, Paz-Ares L, Horn L, et al. Nivolumab versus Docetaxel in Advanced Nonsquamous Non-Small-Cell Lung Cancer. N Engl J Med 2015;373:1627–1639.

3. Brahmer J, Reckamp KL, Baas P, et al. Nivolumab versus Docetaxel in Advanced Squamous-Cell Non-Small-Cell Lung Cancer. N Engl J Med 2015;373:123–135.

4. David S. E ttinger DEW, Dara L. Aisner, et al. NCCN Guidelines Version 3.2019. Non-Small Cell Lung cancer. [https://www.Nccn.Org/professionals/physician_gls/pdf/nscl.Pdf]. Accessed 18 January 2019.

5. Reck M, Rodriguez-Abreu D, Robinson AG, et al. Updated Analysis of KEYNOTE-024: Pembrolizumab Versus Platinum-Based Chemotherapy for Advanced Non-Small-Cell Lung Cancer With PD-L1 Tumor Proportion Score of 50% or Greater. J Clin Oncol 2019;37:537–546.

6. Yu H, Boyle TA, Zhou C, et al. PD-L1 Expression in Lung Cancer. J Thorac Oncol 2016;11:964–975.

7. Hirsch FR, McElhinny A, Stanforth D, et al. PD-L1 Immunohistochemistry Assays for Lung Cancer: Results from Phase 1 of the Blueprint PD-L1 IHC Assay Comparison Project. J Thorac Oncol 2017;12:208–222.

8. Ilie M, Hofman V, Dietel M, et al. Assessment of the PD-L1 status by immunohistochemistry: challenges and perspectives for therapeutic strategies in lung cancer patients. Virchows Arch 2016;468:511–525.

9. Lantuejoul S, Damotte D, Hofman V, et al. Programmed death ligand 1 immunohistochemistry in non-small cell lung carcinoma. J Thorac Dis 2019;11:S89–S101.

10. Cooper WA, Russell PA, Cherian M, et al. Intra- and Interobserver Reproducibility Assessment of PD-L1 Biomarker in Non-Small Cell Lung Cancer. Clin Cancer Res 2017;23:4569–4577.

11. Rehman JA, Han G, Carvajal-Hausdorf DE, et al. Quantitative and pathologist-read comparison of the heterogeneity of programmed death-ligand 1 (PD-L1) expression in non-small cell lung cancer. Mod Pathol 2017;30:340–349.

12. Brunnstrom H, Johansson A, Westbom-Fremer S, et al. PD-L1 immunohistochemistry in clinical diagnostics of lung cancer: inter-pathologist variability is higher than assay variability. Mod Pathol 2017;30:1411–1421.

13. Troncone G, Gridelli C. The reproducibility of PD-L1 scoring in lung cancer: can the pathologists do better? Transl Lung Cancer R 2017;6:S74–S77.

14. Hamilton PW, Bankhead P, Wang YH, et al. Digital pathology and image analysis in tissue biomarker research. Methods 2014;70:59–73.

15. Koelzer VH, Sirinukunwattana K, Rittscher J, et al. Precision immunoprofiling by image analysis and artificial intelligence. Virchows Arch 2019;474:511–522.

16. Salto-Tellez M, Maxwell P, Hamilton P. Artificial intelligence-the third revolution in pathology. Histopathology 2019;74:372–376.

17. Litjens G, Sanchez CI, Timofeeva N, et al. Deep learning as a tool for increased accuracy and efficiency of histopathological diagnosis. Sci Rep-Uk 2016;6.

18. Bejnordi BE, Veta M, van Diest PJ, et al. Diagnostic Assessment of Deep Learning Algorithms for Detection of Lymph Node Metastases in Women With Breast Cancer. Jama-J Am Med Assoc 2017;318:2199–2210.

19. Sirinukunwattana K, Ahmed Raza SE, Yee-Wah T, et al. Locality Sensitive Deep Learning for Detection and Classification of Nuclei in Routine Colon Cancer Histology Images. IEEE Trans Med Imaging 2016;35:1196–1206.

20. Xie W NJ, Zisserman A (2018) Microscopy cell counting and detection with fully convolutional regression networks. Comput Meth Biomech Biomed Eng Imaging Visualization 6: 283–292. https://doi.org/10.1080/21681163.2016.1149104.

21. Esteva A, Kuprel B, Novoa RA, et al. Dermatologist-level classification of skin cancer with deep neural networks. Nature 2017;542:115–118.

22. Mobadersany P, Yousefi S, Amgad M, et al. Predicting cancer outcomes from histology and genomics using convolutional networks. P Natl Acad Sci USA 2018;115:E2970–E2979.

23. Qaiser T, Mukherjee A, Reddy PBC, et al. HER2 challenge contest: a detailed assessment of automated HER2 scoring algorithms in whole slide images of breast cancer tissues. Histopathology 2018;72:227–238.

24. Koelzer VH, Gisler A, Hanhart JC, et al. Digital image analysis improves precision of PD-L1 scoring in cutaneous melanoma. Histopathology 2018;73:397–406.

25. Kapil A, Meier A, Zuraw A, et al. Deep Semi Supervised Generative Learning for Automated Tumor Proportion Scoring on NSCLC Tissue Needle Biopsies. Sci Rep 2018;8:17343.

26. Taylor CR, Jadhav AP, Gholap A, et al. A Multi-Institutional Study to Evaluate Automated Whole Slide Scoring of Immunohistochemistry for Assessment of Programmed Death-Ligand 1 (PD-L1) Expression in Non-Small Cell Lung Cancer. Appl Immunohistochem Mol Morphol 2019;27:263–269.

27. Pan S J, Yang Q. A Survey on Transfer Learning[J]. IEEE Transactions on Knowledge and Data Engineering, 2010, 22(10): 1345–1359

28. Lee MC, Kadota K, Buitrago D, et al. Implementing the new IASLC/ATS/ERS classification of lung adenocarcinomas: results from international and Chinese cohorts. J Thorac Dis 2014;6:S568–580.

29. Bankhead, P. et al. (2017). QuPath: Open source software for digital pathology image analysis. Scientific Reports. https://doi.org/10.1038/s41598-017-17204-5

30. Ficarra E, Di Cataldo S, Acquaviva A, et al. Automated Segmentation of Cells With IHC Membrane Staining. Ieee TBio-Med Eng 2011;58:1421–1429.

31. Doyle S, Monaco J, Feldman M, et al. An active learning based classification strategy for the minority class problem: Application to histopathology annotation. Bmc Bioinformatics 2011;12.

32. Parra ER, Behrens C, Rodriguez-Canales J, et al. Image Analysis-based Assessment of PD-L1 and Tumor-Associated Immune Cells Density Supports Distinct Intratumoral Microenvironment Groups in Non-small Cell Lung Carcinoma Patients. Clinical Cancer Research 2016;22:6278–6289.

33. Catacchio I, Scattone A, Silvestris N, et al. Immune Prophets of Lung Cancer: The Prognostic and Predictive Landscape of Cellular and Molecular Immune Markers. Transl Oncol 2018;11:825–835.

34. Parra ER, Villalobos P, Mino B, et al. Comparison of Different Antibody Clones for Immunohistochemistry Detection of Programmed Cell Death Ligand 1 (PD-L1) on Non-Small Cell Lung Carcinoma. Appl Immunohistochem Mol Morphol 2018;26:83–93.

35. Scheel AH, Dietel M, Heukamp LC, et al. Harmonized PD-L1 immunohistochemistry for pulmonary squamous-cell and adenocarcinomas. Modern Pathol 2016;29:1165–1172.

