## Supplemental figures and tables for "Deep learning approach for automated cancer detection and tumor proportion score estimation of PD-L1 expression in lung adenocarcinoma"

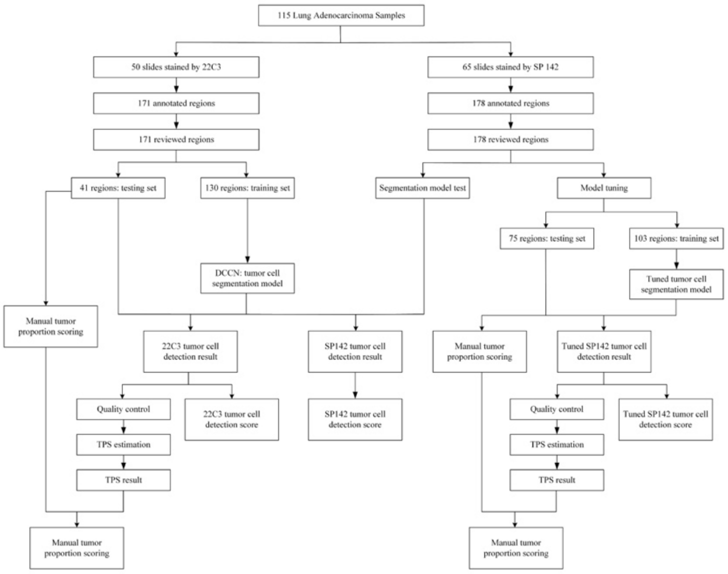


Fig.S1 The workflow of our analysis for the procedure of data analysis and results comparison

Table.S1 The information of patients, including gender, age and cancer stages.

| Clone | | 22C3 pharmDx Assay | SP142 Assay |
|---|---|---|---|
| **Source** | | Dako | ZSGB-BIO |
| **Gender (male: female)** | | 18:25 | 20:27 |
| **Age (average, standard deviation)** | | 56.73±9.69 | 59.33±12.03 |
| **Case of patients** | | 43 | 47 |
| **Cancer Stage** | **1** | 24 | 19 |
| | **2** | 13 | 10 |
| | **3** | 6 | 16 |
| | **4** | 0 | 2 |
| **Regions selected** | | 171 | 178 |
| **Training set** | | 130 | 103 |
| **Test set** | | 41 | 75 |

Table.S2 The concordance of pathologists' labeling. Three random samples were chosen and three pathologists' results were compared here. IoU and concordance matrix between each pair of pathologists were calculated to show the consistence among pathologists' results.

| Sample ID | IoU | Concordance between pairs | | |
|---|---|---|---|---|
| **I** | 0.947 | 0.962 | 0.964 | 0.976 |
| **II** | 0.940 | 0.939 | 0.910 | 0.942 |
| **III** | 0.948 | 0.947 | 0.959 | 0.976 |

Table.S3 Fleiss' and Cohen's Kappa for the agreement among pathologists and between pathologists and AI model. Since we only have two pathologists for SP142 dataset, the median and mean data for manual results are the same with each other, which lead to the same calculated parameters for AI vs. pathologists' median and AI vs. pathologists' mean for SP142 dataset.

| | 22C3 | | SP142 | |
|---|---|---|---|---|
| | **1% cutoff** | **50% cutoff** | **1% cutoff** | **50% cutoff** |
| **Pathologists** | 0.401 | 0.841 | 0.557 | 1.000 |
| **AI vs. pathologists' median** | 0.445 | 0.847 | 0.519 | 0.959 |
| **AI vs. pathologists' mean** | 0.302 | 0.919 | 0.519 | 0.959 |

Table.S4 The average time consumed for image review of both pathologists and AI method.

| Average time/min | | 22C3 | SP142 |
|---|---|---|---|
| Pathologists | W S | 2.08 | 2.42 |
| | J W | 2.98 | 2.91 |
| | Y Z | 3.12 | 3.17 |
| | H W | 2.54 | 2.71 |
| | X L | 2.50 | 2.97 |
| AI method | | About 1.5 | |
